# Transcriptomic Response of Brain Tissue to Focused Ultrasound-Mediated Blood-Brain Barrier Disruption Depends Strongly on Anesthesia

**DOI:** 10.1101/2020.07.24.211136

**Authors:** A.S. Mathew, C.M. Gorick, E.A. Thim, W.J. Garrison, A.L. Klibanov, G.W. Miller, N.D. Sheybani, R.J. Price

## Abstract

Focused ultrasound (FUS) mediated blood brain barrier disruption (BBBD) is a promising strategy for the targeted delivery of systemically-administered therapeutics to the central nervous system (CNS). Pre-clinical investigations of BBBD have been performed on different anesthetic backgrounds; however, the potential influence of the choice of anesthetic on the molecular response to BBBD is unknown, despite its potential to critically affect interpretation of experimental therapeutic outcomes. Here, using bulk RNA sequencing approaches, we comprehensively examined the transcriptomic response of both normal brain tissue and brain tissue exposed to FUS-induced BBBD in mice anesthetized with either isoflurane with medical air (Iso) or ketamine/dexmedetomidine (KD). In normal murine brain tissue, Iso alone elicited minimal differential gene expression (DGE) and repressed pathways associated with neuronal signaling. KD alone, however, led to massive DGE and enrichment of pathways associated with protein synthesis. In brain tissue exposed to BBBD (1 MHz, 0.5 Hz pulse repetition frequency, 0.4 MPa peak-negative pressure), we systematically evaluated the relative effects of anesthesia, microbubbles, and FUS on the transcriptome. Of particular interest, we observed that gene sets associated with sterile inflammatory responses and cell-cell junctional activity were induced by BBBD, regardless of the choice of anesthesia. Meanwhile, gene sets associated with metabolism, platelet activity, tissue repair, and signaling pathways, were differentially affected by BBBD, with a strong dependence on the anesthetic. We conclude that the underlying transcriptomic response to FUS-mediated BBBD may be powerfully influenced by anesthesia. These findings raise considerations for the translation of FUS-BBBD delivery approaches that impact, in particular, metabolism, tissue repair, and intracellular signaling.

## Introduction

The blood-brain barrier (BBB) is essential to maintaining homeostasis in the central nervous system (CNS). The BBB describes a specialized vasculature, consisting of nonfenestrated endothelium, pericytes, astrocytic processes, microglia, and basement membrane working in concert to precisely permit nutrient transport while protecting against toxins and pathogens. However, the efficacy of the BBB also presents a significant neuropharmacological obstacle, preventing 98% of small-molecule therapeutics and nearly 100% of large-molecule therapeutics from accessing the CNS [1]. Significant efforts have focused on strategies to bypass or disrupt the BBB. Methods to bypass the BBB, including intracranial injection and intracerebroventricular infusion, require surgical intervention and thus carry significant risk. Chemical methods to disrupt the BBB, such as mannitol, cause global BBB disruption and lead to considerable neurotoxicity.

Focused ultrasound (FUS) following IV infusion of microbubbles (MB) is a promising approach for BBB disruption (BBBD) [2–4]. In this technique, sound waves produced extracorporeally by an MRI-guided transducer pass through the skull and cause MB circulating in a targeted region of the brain to oscillate. These oscillations produce intravascular cavitation forces capable of disrupting BBB tight junctions and enhancing transport of molecules into the brain parenchyma. FUS induced BBBD is an attractive alternative to surgical and chemical methods as it is targeted, non-invasive, and repeatable. Many therapies normally restricted by the BBB have been successfully delivered with FUS + MB, including antibodies [5–7], chemotherapeutics [8–10], neural stem cells [11,12], and genes [13–15].

BBBD with FUS is reversible and may be applied in a manner that yields little to no histological damage after repeated treatment [3,16,17]. However, recent molecular profiling studies have demonstrated that FUS induced BBBD leads to increased expression of pro-inflammatory cytokines, homing receptors, and damage associated molecular patterns (DAMPs) as well as increased systemic macrophage accumulation in the CNS [18]. These findings are consistent with sterile inflammation (SI), an innate immune response. The potential for FUS to induce local SI has sparked discussion of the cellular implications of FUS, both where additional inflammation may be desirable (such as cancer or Alzheimer’s) or undesirable (such as multiple sclerosis or stroke) [19–22]. Transcriptomic studies have shown that FUS induced SI is proportional to both microbubble dose and FUS acoustic pressure [23,24]. At pressures capable of reliably opening BBB, as measured by MR contrast enhancement, we observed upregulation of proinflammatory transcripts (such as Ccl3, Ccl12, Ccl4, and GFAP) and pathways at 6 h post-FUS, trending toward resolution at 24 h post-FUS, consistent with previous studies [18,24,25]. Recent work has demonstrated the extent of post-FUS SI can be modulated by administration of dexamethasone[26]. Still, knowledge of the contributions of FUS experimental parameters to the SI response as well as non-inflammatory effects on the brain parenchyma remain limited.

One such parameter is general anesthesia. Anesthetic protocols, ubiquitous in preclinical FUS BBBD studies, have been shown to distinctly impact the circulation time of MB and the extent of FUS-induced vascular damage [27,28]. Common anesthetics vary widely in their effects on the CNS, differentially affecting cerebral vasculature, neuronal signaling, inflammation, and metabolism [29–31]. Indeed, a review of the FUS BBBD literature performed by our group (**Table S1**) highlights considerable diversity in anesthetic protocols used in pre-clinical studies of experimental therapeutic efficacy, with isoflurane and ketamine being the most commonly chosen agents. We hypothesize that anesthetics differentially alter the underlying reactivity of the brain parenchyma when FUS is applied, which may produce anesthesia-dependent synergies and conflicts with respect to SI, drug metabolism, or neuronal damage. Herein, we test this hypothesis by detailing the cumulative transcriptome level and pathway level impacts of anesthesia, MB, and FUS on the brain parenchyma.

## Results

### Characterization of FUS-Induced BBBD and Passive Cavitation Analysis

Mice were anesthetized with either isoflurane in medical air (Iso) or ketamine/dexmedetomidine (KD) and treated with Magnetic Resonance-guided Focused Ultrasound (MRgFUS) targeted to the right or left striatum. To assess the extent and localization of BBBD, contrast-enhanced images were collected before and after treatment (**Figure 1A**). The magnitude of signal enhancement was significantly greater in mice anesthetized with Iso compared to KD with respect to fold difference (**Figure 1B**) in mean grayscale intensity in treated vs untreated hemispheres. To evaluate differences in oscillatory activity of circulating MB in response to FUS, we analyzed acoustic emissions data obtained from a listening hydrophone embedded in the therapeutic transducer. Steady oscillation of MB, called stable cavitation, imparts the mechanical forces on vessel walls needed to disrupt the BBB and produces concomitant peaks at harmonics (2f, 3f, 4f, f = operating frequency of the treatment transducer). Meanwhile, unstable oscillation and violent collapse of MB, called inertial cavitation, can produce concomitant broadband signal (in-between harmonics) in the Fourier domain. No significant differences in stable cavitation (as measured by 2^nd^, 3^rd^, 4^th^ harmonics) or inertial cavitation (broadband emission up to 10 MHz) were found between Iso and KD (**Figure 1C**).

**Figure 1:**
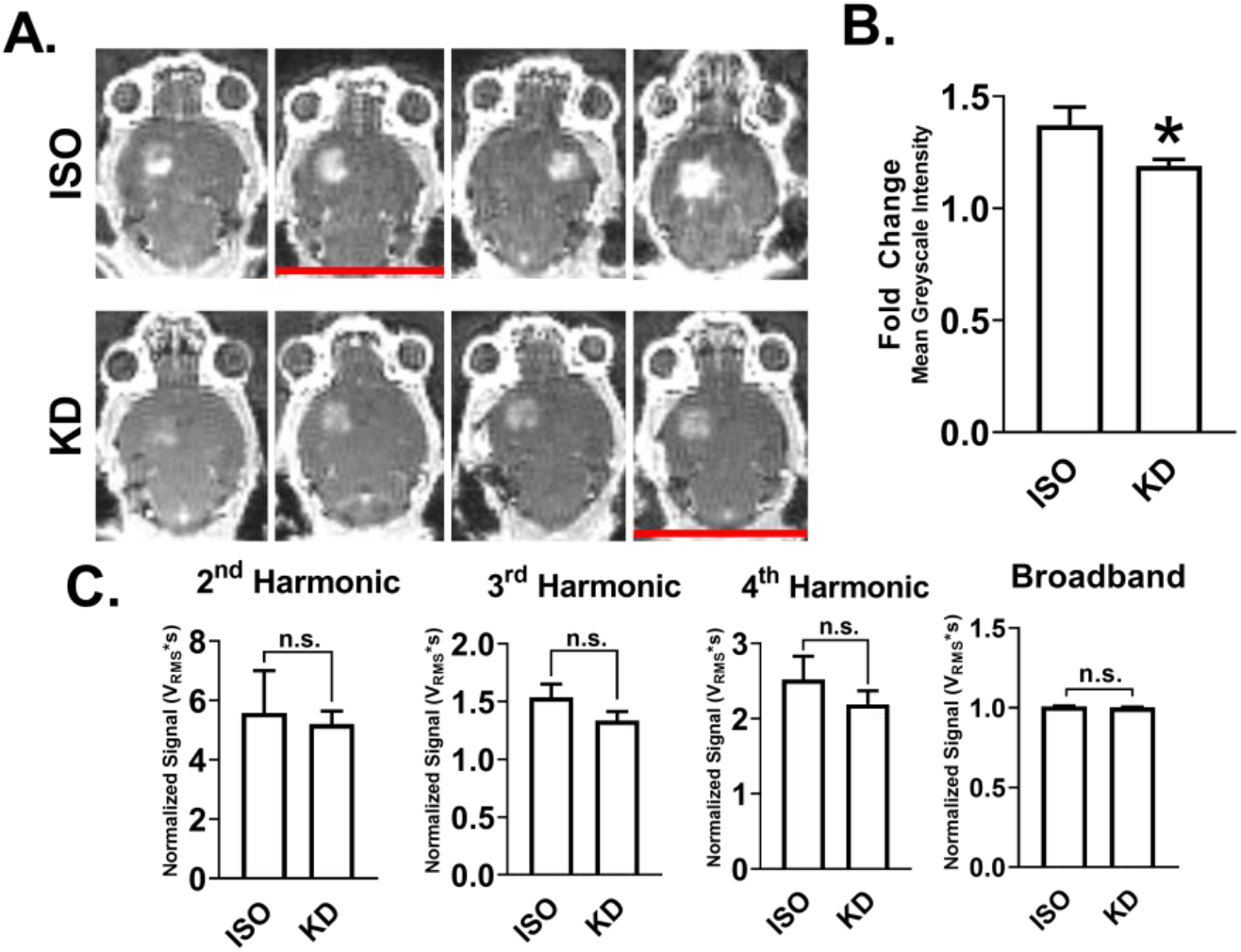
Characterization of FUS-Induced BBBD and Passive Cavitation Analysis. (A) T1-weighted contrast-enhanced 3T MRI images of naïve brains immediately following BBB disruption with FUS+MB. Red lines denote mice that were removed from RNA sequencing analysis due to low RNA integrity number (RIN). (B) Fold difference in mean grayscale signal intensity in contrast-enhanced images in FUS-treated hemisphere relative to contralateral hemisphere. Data are represented as mean with SEM. *p<0.05 (p = 0.0286) by Mann-Whitney test. n=4 mice per group. (C) Acoustic emissions signals (2nd, 3rd, 4th harmonics and broadband) at 0.4 MPa FUS + MB exposure, normalized to 0.005 MPa signal without MB. Data are represented as mean with SEM. No significance was detected by Mann-Whitney test. n=4 mice per group.

### Transcriptomic Variation is Driven Primarily by KD and Secondarily by FUS BBBD

Bulk RNA sequencing was performed on mRNA extracted 6 h post-FUS from the treated region of each brain shown in **Figure 1**. Brains extracted from naïve mice, mice treated with each anesthetic alone, and mice treated with each anesthetic and microbubbles were also sequenced 6 h after treatment. After read alignment and QC, principal components analysis (PCA) was performed on transformed transcript counts from each sample to assess global differences between treatment conditions (**Figure 2A**). Interestingly, the first principal component segregated samples by whether they received KD, with Iso-treated mice clustering more closely to the naïve controls. FUS-treated mice formed a distinct cluster only in the KD treated mice. Similar results were obtained when hierarchical clustering was performed on inter-sample Euclidian distances computed between samples based on their transcript counts (**Figure 2B**). With the exception of one sample, the first branch point of the dendrogram separated samples by KD status, while the second and third branch points distinguished samples by FUS treatment.

**Figure 2:**
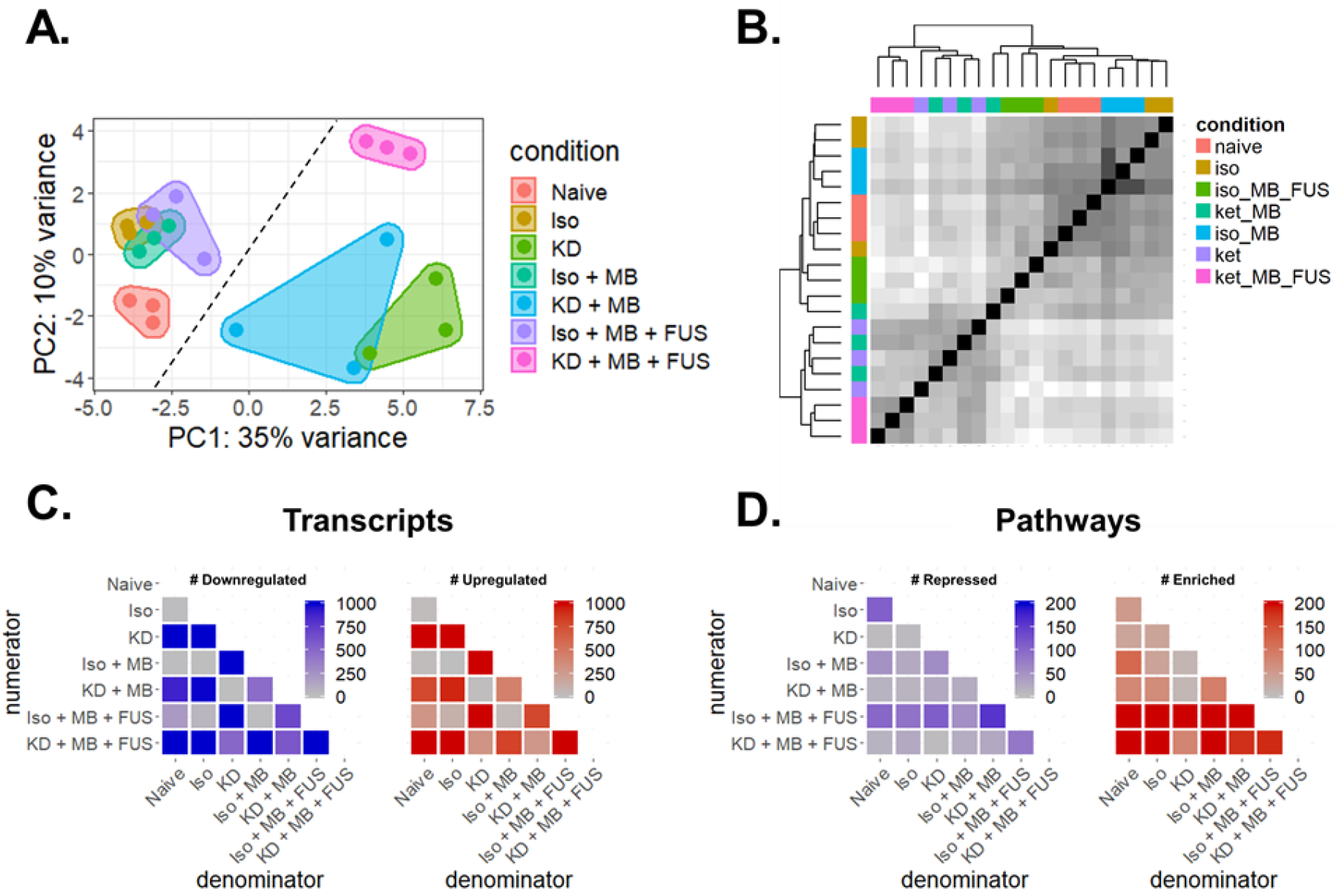
RNA sequencing overview. (A) Principal components analysis of RNA-seq transcript counts after variance stabilizing transformation. Each dot represents a single sample (n = 3 per group). The dashed line separates KD-samples (left of line) from KD+ samples (right of line). (B) Pairwise sample Euclidean distance matrix computed on transcript counts. Each row and column represents a single sample. Hierarchical clustering was performed using complete linkage. Darker shade corresponds to increasing transcriptome similarity. (C) Number of significantly downregulated (left) and upregulated genes (right) for all 21 contrasts of the 7 conditions tested. Each row represents a numerator condition and each column represents a denominator condition. (D) Magnitude of significantly repressed (left) and enriched pathways (right) for all 21 contrasts of the 7 conditions tested. Each row represents a numerator condition and each column represents a denominator condition. For all genes and pathways, significance is defined as p-adjusted < 0.05.

### Overview of Differential Gene Expression and Gene Set Enrichment Analyses

To evaluate relative transcriptomic differences between conditions, differential gene expression contrasts were computed for all 21 unique combinations of the 7 conditions evaluated (**Figure 2C**). KD alone produced the most profound effect on the transcriptome, with over 3000 genes significantly differentially regulated (p-adjusted < 0.05) compared to naïve brain. Regardless of the anesthetic background, FUS and MB produced moderate (on the order of hundreds of differentially expressed genes) and negligible (< 9 differentially expressed genes) effects on gene expression respectively. Iso alone had a marginal effect on the transcriptome, only significantly changing the expression of 26 genes. Next, we performed gene set enrichment analysis (GSEA) to identify biological processes consistent with genes differentially expressed within each contrast (**Figure 2D**). GSEA was performed using the Gene Ontology (GO) Biological Pathways database, wherein each “GO” term represents a collection of genes associated with a particular biological phenomenon. Surprisingly, Iso alone affected more biological pathways than KD, despite KD affecting considerably more genes. The addition of MB changed relatively few biological pathways. FUS had the strongest effect on biological pathways on both anesthetic backgrounds, inducing more pathways than it repressed.

### Anesthetics Differentially Affect the Transcriptome of Normal Brain Tissue

The relative transcriptional impact of Iso and KD on the mouse striatum was marked, with Iso significantly changing expression of 26 genes compared to the 3,291 significantly changed by KD (**Figure 3A**). Iso alone induced a traditional anesthetic transcriptional program of repression of neuronal activity (**Figure 3B**). KD, however, had a minimal effect on these pathways, instead enriching for steps of protein synthesis and targeting (**Figure 3C**). These trends persisted upon addition of MB or FUS. To assess the effect of anesthesia on neuroinflammation, we examined GO processes related to inflammation differentially changed by Iso or KD alone (**Figure 3D**). Both anesthetics induced enrichment of the CCR Chemokine Receptor Binding pathway while only Iso induced the Leukocyte Migration pathway. Interestingly, addition of MB led to loss of significance in CCR Chemokine Receptor Binding enrichment for both anesthetics, while addition of FUS+MB led to further activation of both inflammatory pathways. Iso alone also had a unique effect on development pathways, downregulating neuronal development (likely due to repressing neuronal signaling) and upregulating development of glial cells, oligodendrocytes, and vasculature (**Figure 3E**). In general, addition of MB or MB+FUS led to loss of significance of these pathways. To identify which transcripts contributed to the enrichment or repression of particular pathways, we performed leading edge analysis (LEA). Pecam1 (CD31) was identified as the most significant gene driving the enrichment of the CCR Chemokine Receptor Binding, Leukocyte Migration, and Vasculature Development pathways. Indeed, Pecam1 is one of the few genes induced by Iso with an adjusted p-value less than 0.05.

**Figure 3:**
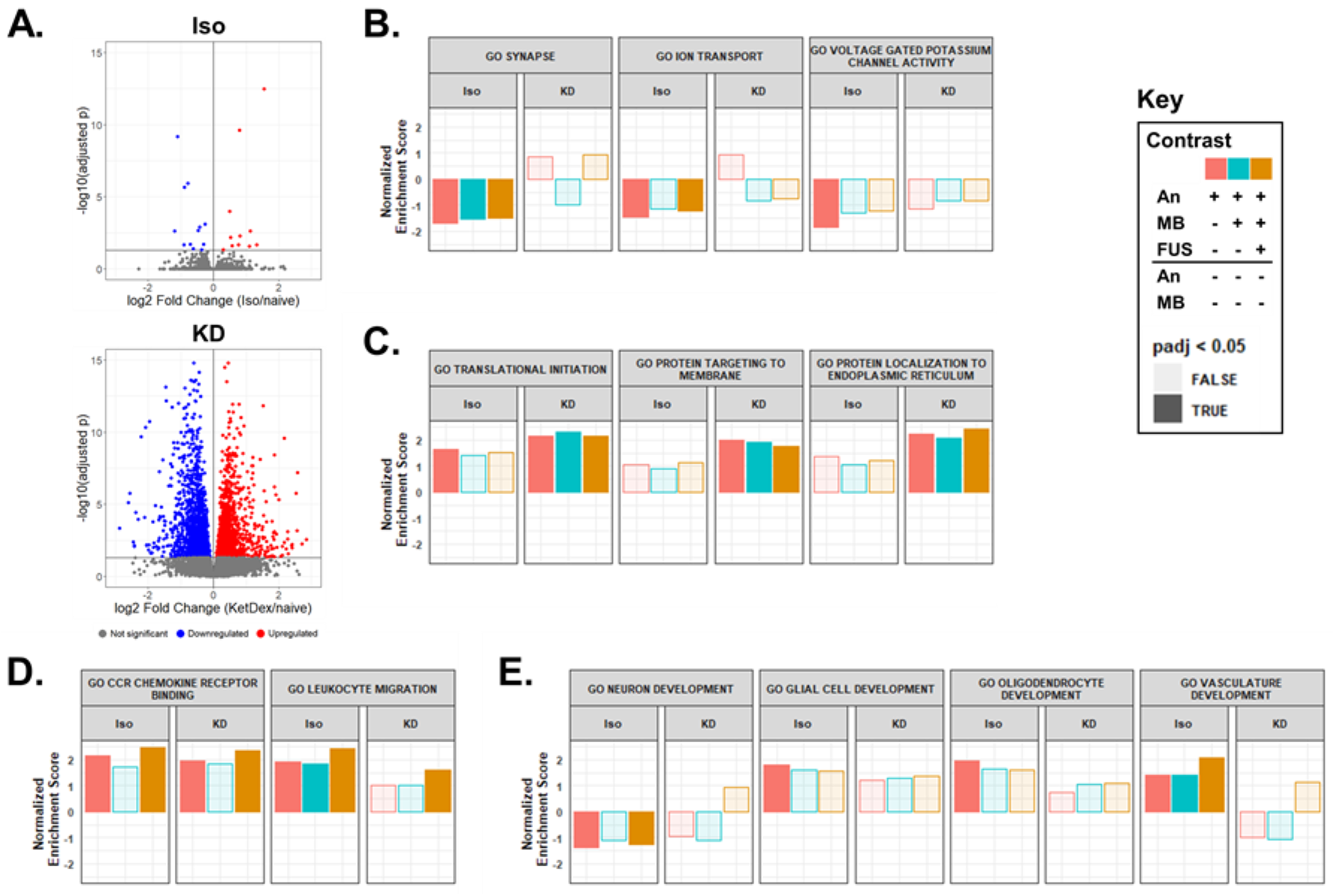
Anesthetics Differentially Affect the Transcriptome of Normal Brain Tissue. (A) Volcano plots of differentially regulated transcripts 6 h post anesthesia delivery with Iso (top) or KD (bottom) compared to naïve controls. (B-E) Normalized Enrichment scores (NES) for gene sets associated with (B) neuronal signaling, (C) protein synthesis, (D) inflammation, and (E) development. GSEA was computed based on ranked DGE from (An, red), An+MB (blue), and An+MB+FUS (gold) against naïve controls for Iso and KD. Opaque bars indicate an adjusted p-value < 0.05.

### Anesthetics Differentially Affect the Transcriptome of Brain Tissue Exposed to FUS BBBD

We next sought to compare gene expression changes induced by FUS BBBD when performed under Iso (Iso-FUS) vs KD (KD-FUS). First, we evaluated the extent and overlap of differentially expressed genes (**Figure 4A**) and differentially regulated pathways (**Figure 4B**), controlling for changes due to anesthesia + MB alone. While more genes were differentially regulated by KD-FUS, more gene sets were significantly enriched/repressed by Iso-FUS. Interestingly, despite minimal intersection of transcript identities between the two BBBD conditions, 41% of the pathways significantly induced by KD-FUS were also significantly induced by Iso-FUS. Second, we identified 6 categories of biological pathways consistently changed by Iso-FUS, KD-FUS, or both (**Figure 4C**). Regardless of the anesthetic background, FUS led to enrichment of genes involved in endothelial cell activity, including pathways associated with cell-cell adhesion and angiogenesis. Iso-FUS induced these pathways more significantly, and additionally led to the expression of genes associated with leukocyte adhesion. Similarly, both FUS conditions led to activation of many inflammation pathways, with the breadth and depth of these responses substantially enhanced in the Iso-FUS condition. Notably, the MHC class I and MHC class II antigen processing and presentation pathways were only upregulated when comparing KD-FUS treated mice to naïve controls. We found the most significant divergence between Iso-FUS and KD-FUS when comparing metabolic pathways. Iso-FUS led to repression of broad and specific metabolic programs while several of these were enriched by KD-FUS. Consistent with significant inflammation and endothelial activation, platelet activity was enhanced by Iso-FUS, while these pathways were relatively unchanged by KD-FUS. Gene sets associated with tissue repair were enriched by FUS under both anesthetics and those associated with neurogenesis were additionally upregulated by KD-FUS only. Signaling pathways engaged by FUS treatment independent of anesthesia included VEGFR signaling, Wnt signaling, and the NF-κB signaling pathway. STAT, SAPK, dopamine, and integrin signaling were further enriched only in Iso-FUS contrasts.

**Figure 4:**
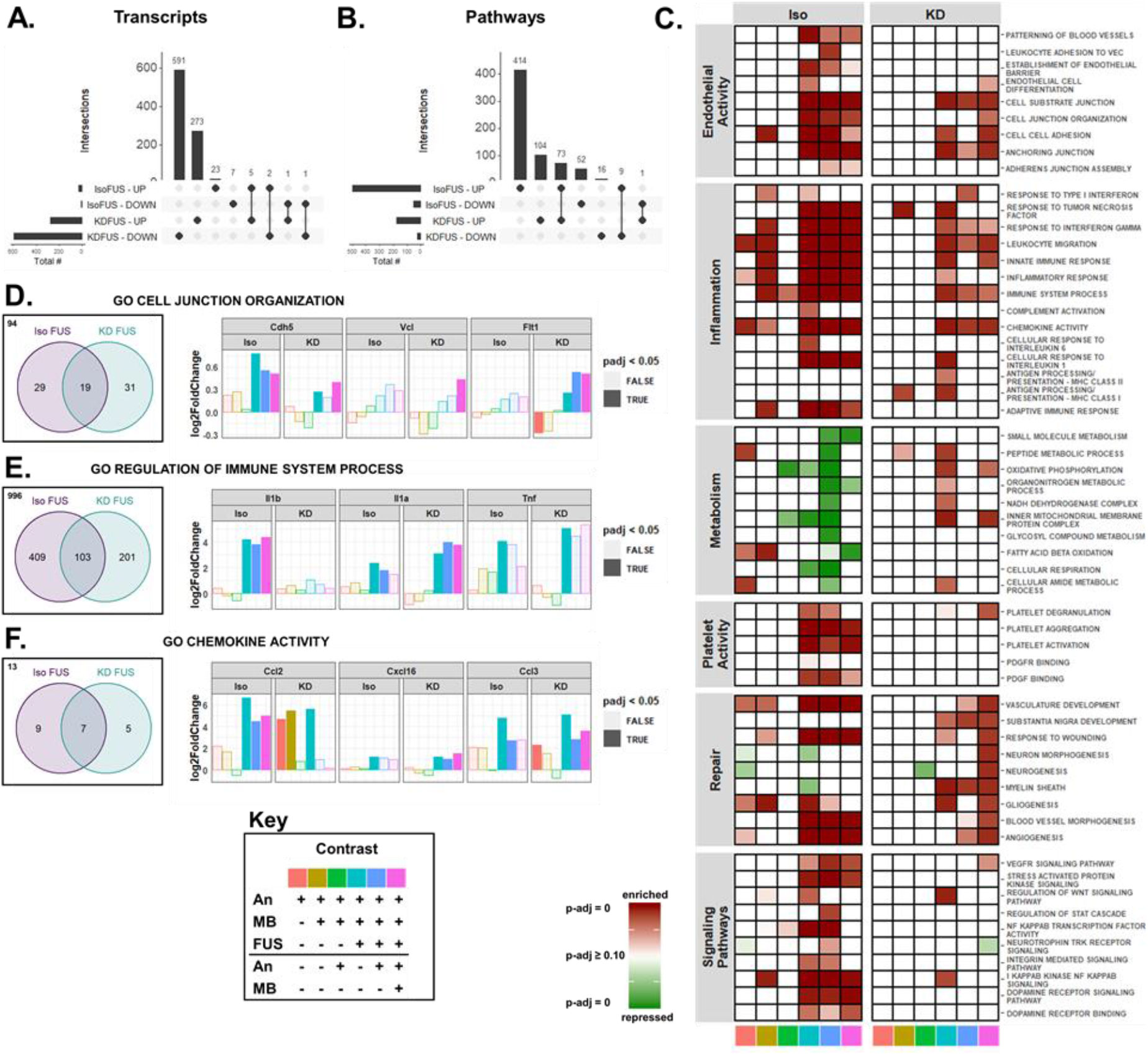
Anesthetics Differentially Affect the Transcriptomic of Brain Tissue Exposed to FUS BBBD. UpSetR plots for evaluating intersections of upregulated and downregulated transcripts between IsoFUS and KDFUS, controlling for the effects of anesthesia and MB. (B) UpSetR plots for evaluating intersections of enriched and repressed pathways between IsoFUS and KDFUS, controlling for the effects of anesthesia and MB alone. (C) Heatmap showing significance of repression (green) or enrichment (red) of pathways (rows) associated with endothelial activity, inflammation, metabolism, platelet activity, repair, and signaling for multiple contrasts (columns), separated by anesthetic. Contrast identities are shown by the color at the bottom of the column, corresponding to the key. Full opacity corresponds to an adjusted-p-value of 0, while full transparency corresponds to an adjusted p-value ≥ 0.10. (D-F) Venn diagrams (left) of leading edge transcripts and selected leading edge transcript expression (right) for (D) Cell Junction Organization (GO:0034330), (E) Regulation of Immune Process (GO:0002682), and (F) Chemokine Activity (GO:0008009) gene sets, separated by anesthetic background. Bar color represents the contrast, corresponding to the key. Opaque bars indicate an adjusted p-value < 0.05. Each color in the key corresponds to a specific pairwise comparison of Anesthesia (An), An + MB, and An + MB + FUS for either Iso or KD, specifying the numerator (above the black line), and denominator (below the black line). For example, pink corresponds to the ratio of gene expression for mice treated with An + MB + FUS to those treated with just An + MB.

To further compare the effect of anesthesia on FUS BBBD, we performed leading edge analysis (LEA) on selected gene sets enriched by both Iso-FUS and KD-FUS. Comparing transcripts in the LEA of the (Iso + MB + FUS)/(Iso + MB) contrast against those in LEA of the (KD + MB + FUS)/(KD + MB) contrast for the same pathway allows us to address whether FUS is achieving the same “end” (pathway enrichment) by similar “means” (transcript regulation) on different anesthetic backgrounds. We performed comparative LEA on gene sets associated with cell-cell junctions and inflammation, as these were the most consistently induced by both Iso-FUS and KD-FUS. Out of the 173 genes in the Cell Junction Organization gene set (GO:0034330), Iso-FUS and KD-FUS enriched 48 and 50 respectively (**Figure 4D**). 19 transcripts were found in the leading edge of both anesthetics including Cdh5 (VE-Cadherin), Vcl, and Flt1. While all 3 of these transcripts were significantly upregulated by KD-FUS across multiple contrasts, only Cdh5 was significantly upregulated by FUS under Iso. Notably, when compared to naïve controls alone, KD alone significantly downregulated Flt1 and KD + MB led to a trending decrease (p-adj = 0.06). We next compared the LEA overlap on the Immune System Process gene set (GO:0002682), a broad collection of 1709 genes associated with the immune system (**Figure 4E**). Iso-FUS and KD-FUS enriched 512 and 304 of these respectively, with 103 genes enriched by both. IL-1α was found in both LEAs and significantly upregulated across multiple contrasts while IL-1β was only found in the Iso-FUS LEA and indeed only significantly upregulated in Iso-only FUS contrasts. TNFα was found in both LEAs to be significantly upregulated by FUS under both anesthetics when compared to naïve controls, and trending upward in other FUS contrasts. To narrow the scope of immune system-related LEA overlaps, we repeated this analysis on the Chemokine Activity gene set (GO:0008009) which only contains 34 genes (**Figure 4F**). Iso-FUS and KD-FUS enriched 16 and 12 chemokines respectively, 7 of which were shared. Iso-FUS induced the strongest Ccl2 upregulation regardless of the control condition. KD alone induced a comparable upregulation of Ccl2 with no additional effect due to FUS. Cxcl16 however was more strongly induced with KD-FUS than Iso-FUS when controlling for anesthetic. Ccl3 was upregulated by FUS under both anesthetics as well as KD alone. In summary, while FUS promotes phenotypes such as cell junction organization, inflammation, and chemokine activity independent of anesthetic, the nature of the transcripts mediating these effects are often anesthesia-dependent.

### Anesthetics Differentially Affect Transcripts Associated with BBB Structure and Function

We next evaluated the effects of anesthesia, MB, and FUS on transcripts known to be associated with the BBB [32]. Iso-FUS upregulated transcripts mediating leukocyte adhesion, including E-selectin, P-selectin, and Icam1 (**Figure 5A**). Icam1 was also upregulated by KD alone when compared to sham and by KD-FUS when compared to KD or KD + MB. With respect to BBB tight junction transcripts, FUS upregulated Cldn5 and Emp1 independent of anesthetic (**Figure 5B**). KD alone led to downregulation of Ocln and Tjp1. We next evaluated the effect of our experimental conditions on BBB transporter transcripts and observed heterogeneous effects (**Figure 5C**). In general, KD led to significantly more DGE in this category than Iso, with very few transcripts changing their expression due to FUS or MB on either anesthetic background. This trend was even more extreme when evaluating BBB transcripts involved in transcytosis and other miscellaneous functions (**Figure 5D**); KD was the only variable significantly changing the expression of transcripts in this class.

**Figure 5:**
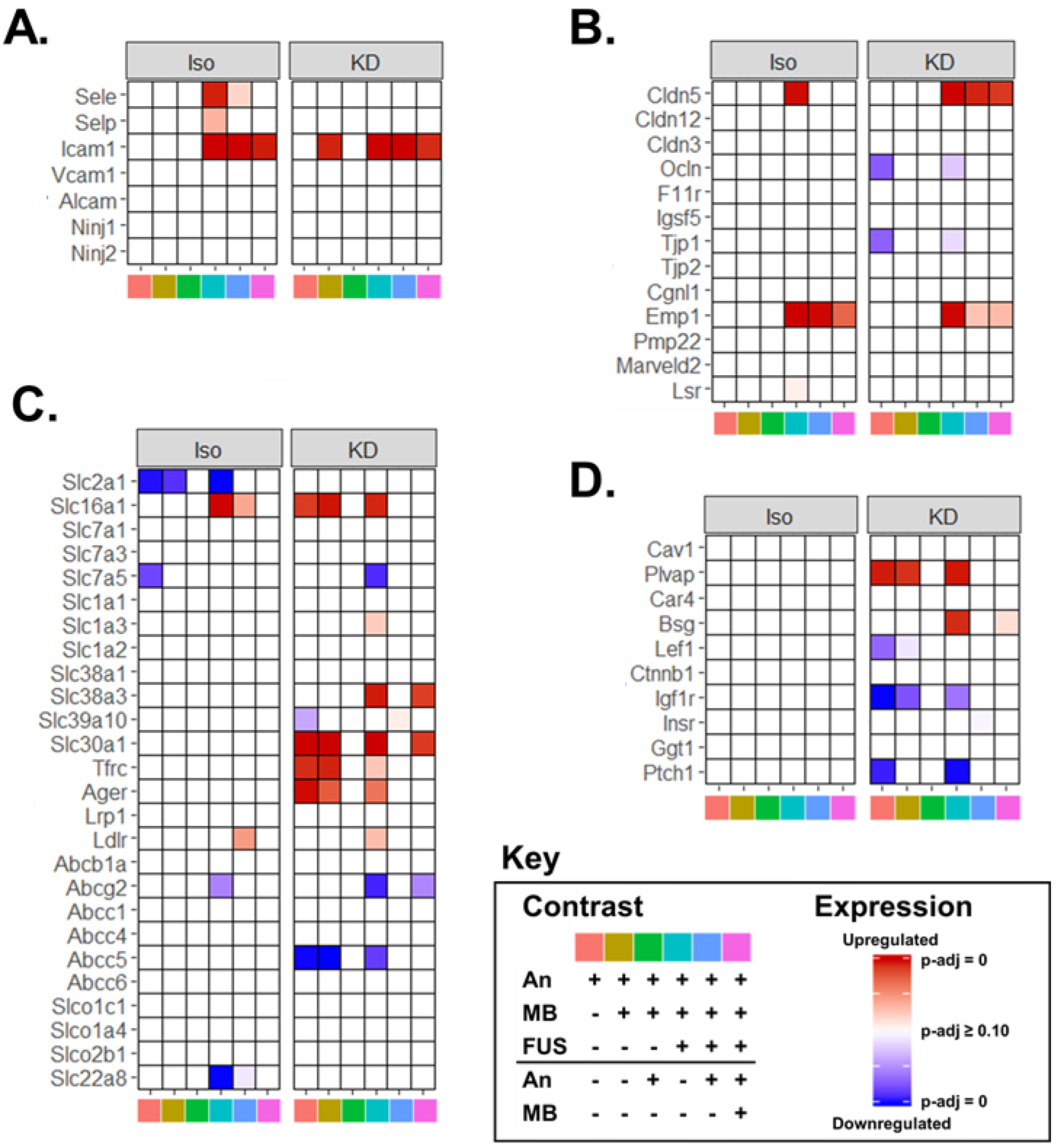
Anesthetics Differentially Affect Transcripts Associated with BBB Structure and Function. (A-D) Heatmaps of significance of upregulation (red) or downregulation (blue) for selected genes (rows) across multiple contrasts (columns), separated by anesthetic for transcripts associated with BBB structure and function. Selected categories include (A) leukocyte adhesion, (B) BBB tight junctions, (C) transporters, and (D) transcytosis/miscellaneous. Contrast identities are shown by the color at the bottom of the column, corresponding to the key. Full opacity corresponds to an adjusted-p-value of 0, while full transparency corresponds to an adjusted p-value ≥ 0.10. Each color in the key corresponds to a specific pairwise comparison of Anesthesia (An), An + MB, and An + MB + FUS for either Iso or KD, specifying the numerator (above the black line), and denominator (below the black line). For example, pink corresponds to the ratio of gene expression for mice treated with An + MB + FUS to those treated with just An + MB.

### Tissue Damage Elicited by FUS BBBD is Minimal and Not Affected by Anesthetic

Given the anesthesia-dependence of BBBD and FUS-induced gene expression, we next tested whether anesthesia significantly affected the extent of damage in the brain parenchyma after treatment with the same FUS pressure. To address this, we performed histological analysis of murine brains treated with combinations of Iso, KD, and FUS (**Figures 6A-D**). Brains treated with 0.8 MPa (twice the acoustic pressure of our standard BBBD protocol) were used as positive controls for damage. We scored multiple transverse sections from each condition for RBC extravasation and vacuolation (**Figure 6E**). With the exception of the 0.8 MPa positive control group, all conditions tested elicited minimal evidence of damage. Thus, we confirmed that BBBD using the FUS parameters selected here elicits little to no histological damage, independent of whether Iso or KD is used.

**Figure 6:**
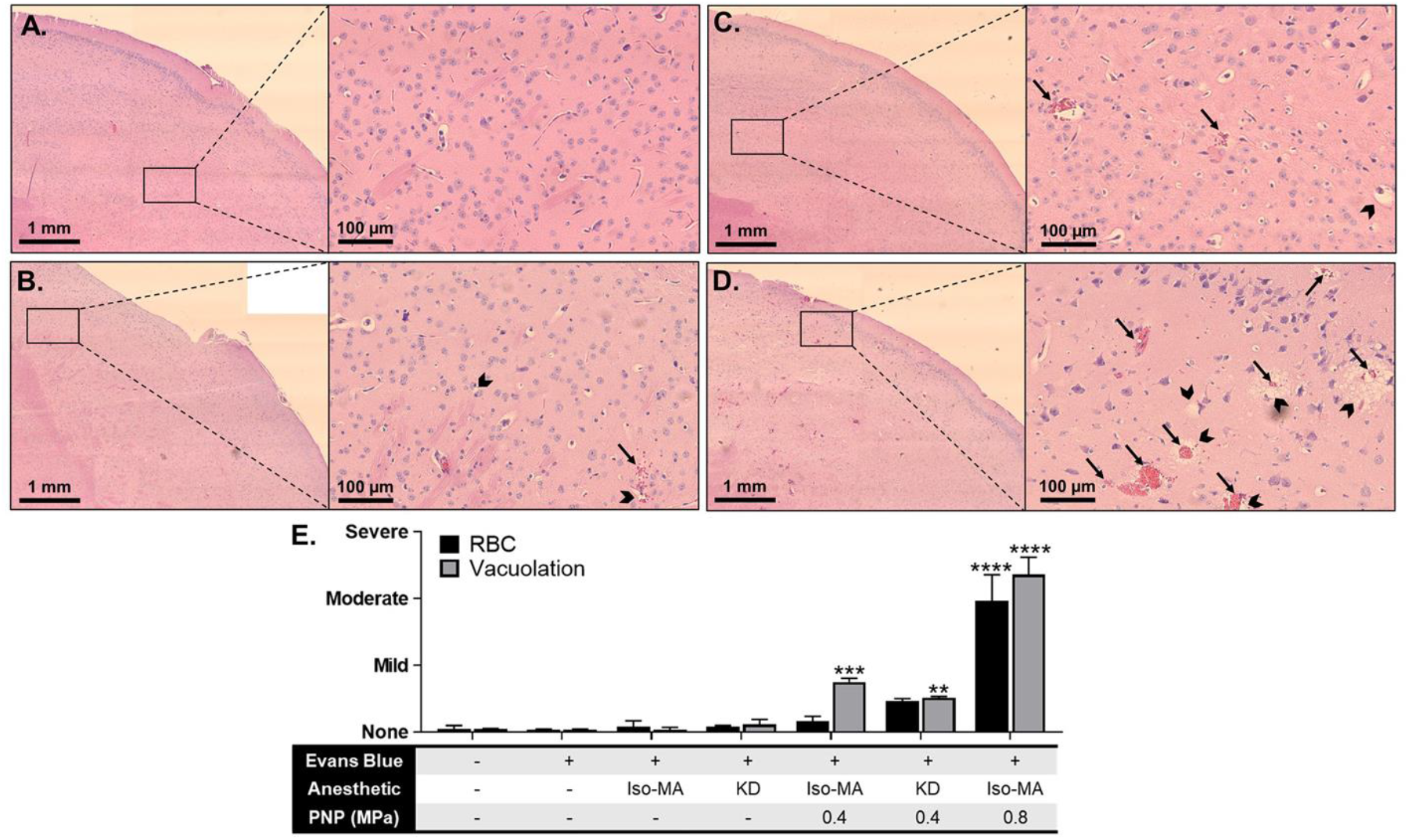
Tissue Damage Elicited by FUS BBBD is Minimal and Not Affected by Anesthetic. Representative 4x stitched (left) and 20x (right) H&E images of murine right striatum either (A) untreated or treated with (B) IsoMA-FUS at 0.4 MPa, (C) KD-FUS at 0.4 MPa, or (D) IsoMA-FUS at 0.8 MPa. Arrows indicate RBC extravasation, chevrons indicate vacuolation. (E) Scoring of RBC extravasation (black bars) and vacuolation (grey bars). Data are represented as mean + SEM. **p < 0.01, **p 0 < 0.001, ****p < 0.00001 by one-way ANOVA followed by comparison against naïve with Dunnett’s multiple comparison test.

## Discussion

BBBD mediated by FUS-activated MB has emerged as a promising technique for the image-guided and non-invasive delivery of therapeutics to the CNS. Though this procedure is safe, our understanding of cellular responses to FUS BBBD at the transcriptional level is still limited. This knowledge gap becomes especially significant when considering that pre-clinical BBBD studies have been performed on a multitude of different anesthetic backgrounds (**Table S1**), a factor that could complicate the interpretation of how experimental therapeutic outcomes will translate to human applications, wherein such anesthetics are not utilized. Our study systematically addressed how choice of general anesthetic shapes acute transcriptomic responses to FUS with respect to sterile inflammation, endothelial activity, metabolism, platelet activity, repair, molecular signaling, and BBB-associated genes. Ultimately, we conclude that the underlying transcriptomic response to FUS-mediated BBBD may be strongly influenced by the choice of anesthetic. Such responses may synergize and/or conflict with responses generated by the therapeutic approach itself. Thus, our results provide a framework for rational anesthesia selection for preclinical BBBD studies and will likely find utility when comparing clinical outcomes to pre-clinical results for FUS mediated BBBD drug and gene delivery approaches.

As shown in **Figure 1**, the magnitude of the FUS BBBD by MR-contrast enhancement depended on anesthetic. Anesthesia-dependent differences in BBBD have been reported previously, in which Ketamine/Xylazine (another a2-adrenergic receptor agonist) led to greater contrast enhancement and histological damage than isoflurane (with O_2_ as the carrier gas) after FUS application [27]. We observed greater contrast enhancement with Iso (with MA, a carrier gas known to lead to longer MB circulation times and BBBD than O_2_) than KD, despite comparable levels of MB cavitation [28]. This finding may be attributable to differential vascular effects of Iso vs. KD. Iso causes vasodilation in the BBB and increases cerebral blood flow (CBF) [33,34]. Ketamine is also thought to cause vasodilation and an increase in CBF [35,36]. Dexmedetomidine, however, produces vasoconstriction and a decrease in CBF [37,38]. In a direct comparison, a 2010 study showed higher CBF with isoflurane (with MA) than Ketamine/Xylazine in rats [39]. Enhanced CBF, given the same degree of endothelial disruption, would lead to enhanced gadolinium accumulation and similar MB cavitation, as we observed in **Figure 1.**Iso alone has been shown to increase BBB permeability and impart BBB structural alterations [40,41]. Analogously, the cerebrovasodilatory agents mannitol or alcohol lead to BBB disruption by cellular shrinkage or augmenting matrix metalloproteinases activity respectively [42,43]. Though we did not test the effect of Iso alone on BBB integrity herein, we postulate direct effects of Iso on the BBB may further potentiate disruption by FUS, leading to the elevated contrast enhancement we observed compared to KD-FUS. The mechanisms by which Iso and KD differentially prime the BBB for disruption by FUS-activated MB may contribute to subsequent differences in gene expression and warrant further investigation.

PCA and hierarchical clustering performed on variance-stabilizing transformed RNA-seq counts data revealed the relative contributions of Iso, KD, MB, and FUS to intersample variability with respect to CNS gene expression (**Figures 2A-B**). The most striking of these was KD, inducing DGE (p-adjusted <.05) of 3291 genes when compared to naïve controls (**Figure 3A**). Whether this profound change in gene expression is attributable to ketamine, dexmedetomidine, or both is unclear. Microarray studies of developing rat brain have shown a similar magnitude of acute differential gene expression from ketamine alone [44]. More specifically, investigators reported 819 differentially expressed genes with fold change >1.4, p-adj < 0.05 compared to the 1182 meeting these criteria in our study at an identical timepoint. Though ketamine’s mechanism of action is still unclear, recent studies into its rapid anti-depressant action suggest ketamine indirectly suppresses eukaryotic elongation factor 2 kinase (eEf2K), leading to increased protein translation [45]. This mechanism is in agreement with our pathway level findings (**Figure 3C**). Though fewer transcriptomic level studies exist for dexmedetomidine, it is known to acutely augment transcriptional programs associated with inflammation and circadian rhythm [46,47]. In stark contrast to KD, we found Iso had a negligible impact on gene expression, only significantly altering the expression of 26 genes. This finding is in close agreement with existing acute transcriptomic studies of inhalable anesthetics in rats, which report between 0 and 20 differentially expressed genes [48,49]. Interestingly, despite weak changes in expression magnitude, Iso changed regulation of significantly more pathways than KD (**Figure 2D**). We thus hypothesize that, while Iso influences more targeted transcriptional programs, the combination of ketamine and dexmedetomidine elicits wide-ranging, complex transcription thereby preventing GSEA from detecting discrete pathway enrichment.

We observed increases in inflammatory signatures elicited by both anesthetics (**Figure 3D**). Of the few genes upregulated by Iso alone, a surprising number were immune-associated. Some examples include upregulation of T-cell associated markers Ly6a and Ctla2a, upregulation of adhesion markers Pecam1 and CD93, and downregulation of Nfkbia, the protein product of which inhibits NF-κB. Indeed, activation of NF-κB has been proposed as a mechanism by which volatile anesthetics elicit neuroinflammation [50,51]. Several rodent studies have demonstrated volatile anesthetics can also acutely induce expression of IL-6, IL-1β, and activated caspase-3 [52–55]. It is worth noting that under conditions of CNS stress, including ischemia or LPS exposure, volatile anesthetics have been shown to attenuate inflammation, suggesting that these drugs may contribute to maintaining homeostasis in the brain, rather than being strictly pro- or anti-inflammatory [56–59]. KD also induced signatures associated with inflammation, though to a lesser extent and with a less clear mechanism than Iso. At the chemokine level, for example, we found KD significantly upregulated Ccl17, Ccl2, Ccl3, and Ccl6 with minor but significant downregulation of Cxcl12 and Cx3cl1. These mixed effects may be caused by contrasting neuroinflammatory effects produced by ketamine and dexmedetomidine. Ketamine has been shown to be acutely inflammatory in naïve mice, increasing levels of IL-6, IL-1β, and TNF-α [60], while Dexmedetomidine tends to protect against neuroinflammation [61–64]. The ability of each anesthetic to amplify or protect against SI induced by FUS may be an important experimental consideration for future preclinical FUS work.

SI caused by FUS-activated MB has raised concerns over its feasibility for repeated clinical application. Studies have demonstrated this response can last for at least 24 h after a single sonication, and is dependent on MB dose and FUS pressure [18,23–25]. Proposed causes for this response include damage due to direct and indirect acoustic forces on the neurovascular unit, ischemia reperfusion injury due to FUS-induced vasospasm, and leakage of blood into the brain parenchyma [18,23–25]. Our unbiased bioinformatics analyses suggest that a confluence of these mechanisms is implicated, and can be affected by choice of anesthetic (**Figure 4C**). Pathways enriched by both Iso-FUS and KD-FUS clearly indicate extensive cytokine production, possibly initiated by damage associated molecular pattern (DAMP) release and pattern recognition receptor (PRR) signaling. In general, Iso-FUS led to more extensive activation of the immune response compared to KD-FUS, enriching signatures associated NF-κB signaling, consistent with previous studies [18,23,25]. However even for pathways with similar enrichment, LEA suggest anesthesia affects the quality of FUS-induced SI. The anesthesia dependent induction of IL-1β and IL-1α provides an excellent example (**Figure 4E**). Though both members of the IL1 family bind to the same receptor, several findings point to fundamentally different upstream triggers and downstream consequences. IL-1α is constitutively expressed and acts as a dual-function cytokine, possessing both intracellular activity as a proinflammatory transcription factor and extracellular activity as a DAMP [65,66]. IL-1β, however, is induced by NOD-, LRR- and pyrin domain-containing protein 3 (NLRP3) inflammasome activation [67]. Importantly, it has been shown that these two cytokines recruit different populations of myeloid cells and represent distinct stages of the SI response [68]. Thus, anesthesia may impact the temporal relationship between FUS application and SI. Enrichment of junctional assembly pathways, VEGF signaling, and angiogenesis supports FUS-induced activation of endothelial cells, leading to both recruitment of leukocytes and barrier repair, especially under Iso. Of note, we observed significant upregulation of claudin-5 transcript, whose tight junction protein product is essential to BBB integrity, in both FUS groups. This may indicate initiation of transcriptional programs to repair the disrupted barrier (**Figure 5B**). In contrast, a microarray study of brain microvessels did not detect significant differences in claudin-5 post-FUS [25]. This discrepancy could be due to differences in species (i.e. mouse vs. rat), the source of the analyzed tissue in the brain, anesthesia protocol, and several focused ultrasound and microbubble parameters. Downregulation of multiple metabolic pathways in Iso-FUS contrasts further suggests Iso may prime the BBB for more significant alteration than KD.

Despite such differential responses at the transcriptional level, FUS applied under both anesthetics led to little to no generation of petechiae by H&E (**Figure 6**). With respect to coagulation signatures by RNA-seq, only Iso-FUS led to increased platelet activity despite no significant difference in RBC extravasation compared to KD-FUS (**Figure 4C**). While Iso has been shown to have minimal effect on platelet activity [69–71], both ketamine and dexmedetomidine have been shown to reduce coagulability [72–75]. We hypothesize that KD thus minimizes the inflammatory response resulting from blood products in the brain parenchyma compared to Iso upon FUS application.

Transient SI can provide beneficial effects in certain disease contexts with respect to clearance and regeneration [76]. Indeed, this may be the primary mechanism by which FUS promotes Aβ plaque clearance in models of Alzheimer’s disease [77]. Similarly, neurogenesis observed after FUS may be attributable to tissue repair mechanisms preceded SI [78,79]. We observed activation of repair mechanisms by FUS, though to different extents depending on the anesthetic chosen. The observation that KD promotes stronger signatures of repair and weaker signatures of inflammation, endothelial activation, coagulation, and metabolic alteration supports its use over Iso for pathologies where further CNS stress is undesirable.

Our investigation has some limitations. First, RNA-sequencing only provides transcript-level information and several studies highlight that mRNA may not always correlate proportionally to protein expression [80–83]. This risk is mitigated at the pathway level, where we present significant alteration of large families of genes consistently up or downregulated by FUS and/or anesthesia. We further assert that the high intragroup consistency along with the absolute magnitude of differential gene or pathway level changes we present make noise an unlikely driver of the diverse changes we observe. However we also note that because RNA-seq was performed on bulk tissue, it is not easy to distinguish changes in transcription from changes in relative cell numbers. Protein and phenotypic studies may provide additional insight into the consequences of the results generated herein. Next, whether transcriptional changes in Iso-FUS mice are a consequence of isoflurane’s interaction with FUS or enhanced BBB permeability is unclear. Finally, not all experiments were performed on the same FUS-system. Though transducer frequencies and acoustic pressures were matched between systems, it is possible that differences in transducer geometries produced confounders in experimental endpoints.

We present here a detailed account of how Iso and KD, the two most commonly used anesthetics in preclinical FUS BBBD studies, differentially affect CNS responses to FUS-activated MB. At the same acoustic pressure, FUS induced similar profiles of MB cavitation and measures of damage regardless of the anesthetic. RNA sequencing performed acutely after treatment with combinations of Iso, KD, MB, and FUS revealed distinct contributions from each. Specifically, while Iso alone produced transcriptomic profiles nearly identical to those of naïve mice, it also elicited stronger signatures of stress in the neurovascular unit when combined with FUS. KD, however, induced sweeping transcriptome changes alone, but blunted markers of SI while promoting gene sets associated with tissue repair upon FUS application compared to Iso-FUS. These results provide important context for previous preclinical FUS studies, and underscore anesthesia as an important experimental variable to consider for future work. More research is required to understand whether the findings described herein are maintained at the protein level and how anesthesia-dependent responses to FUS evolve with varying FUS parameters, MB characteristics, and time.

## Materials and Methods

### Animals

11 week old female C57BL/6 mice were purchased from Jackson and maintained on a 12/12 hour light/dark cycle. Mice weighed between 22 and 28 g and were given food and water *ad libitum*. All animal experiments were approved by the Animal Care and Use Committee at the University of Virginia and conformed to the National Institutes of Health regulations for the use of animals in research.

### Anesthesia

Mice in groups designated “KD” received 50-70 mg/kg Ketamine and 0.25-0.5 mg/kg Dexmedetomidine via intraperitoneal injection with no additional maintenance or reversal drug given. Mice in groups designated as “Iso” or “Iso-MA” were placed in an induction chamber and received isoflurane delivered to effect in concentrations of 2.5% in medical air using a vaporizer. For isoflurane groups, anesthesia was maintained via nosecone for a total of 90 minutes.

### MRgFUS mediated BBBD

Once anesthetized, a tail vein catheter was inserted to permit intravenous injections of MBs and the MRI contrast agent. The heads of the mice were shaved and depilated, and the animals were then placed in a supine position over a degassed water bath coupled to an MR-compatible small animal FUS system (RK-100; FUS Instruments, Toronto, Canada). The entire system was then placed in a 3T MR scanner (Magnetom Prisma; Siemens Medical Solutions, Malvern, Pennsylvania). A 3.5 cm diameter receive RF coil, designed and built in-house, was placed around the head to maximize imaging SNR. Baseline three-dimensional T1-weighted MR images were acquired at 0.3 mm resolution using a short-TR spoiled gradient-echo pulse sequence and used to select 4 FUS target locations in and around the right or left striatum.

Mice received an injection of albumin-shelled MBs (1 × 10^5^ MBs/g b.w.), formulated as previously described [14,84,85]. Sonication began immediately after clearance of the catheter. Sonications (4 spots in a 2×2 grid) were performed at 0.4 MPa peak-negative pressure (PNP) using a 1.1 MHz single element focused transducer (FUS Instruments, Toronto, Canada) operating in 10 ms bursts, 0.5 Hz pulse repetition frequency and 2 minutes total duration. Immediately following the FUS treatment, mice received an intravenous injection of gadolinium-based contrast agent (0.05 ml of 105.8 mg/ml preparation; Multihance; Bracco Diagnostics), and contrast-enhanced images were acquired to assess BBBD using the same T1-weighted pulse sequence mentioned above.

### Passive Cavitation Detection

Acoustic emissions were detected with a 2.5 mm wideband unfocused hydrophone mounted in the center of the transducer. Acoustic signal was captured using a scope card (ATS460, Alazar, Pointe-Claire, Canada) and processed using an in-house built MATLAB (MathWorks) algorithm. Acoustic emissions at the fundamental frequency, harmonics (2f, 3f, 4f), sub harmonic (0.5f), and ultra-harmonics (1.5f, 2.5f, 3.5f) were assessed by first taking the root mean square of the peak spectral amplitude (Vrms) in each frequency band after applying a 200 Hz bandwidth filter, and then summing the product of Vrms and individual sonication duration over the entire treatment period. Broadband emissions were assessed by summing the product of Vrms and individual sonication duration for all remaining emissions over the entire treatment period.

### Bulk RNA Sequencing and Analysis

6 hours after treatment, mice were euthanized via an overdose of pentobarbital sodium and phenytoin sodium. Immediately following euthanasia, the mouse brains were harvested and the front right quadrants were excised (with the exception of 1 mouse, which had FUS treatment on the left), placed in RNAlater (Qiagen), and stored at −80 °C. RNA extraction was performed using the RNeasy Mini Kit (Qiagen). mRNA was isolated using the NEBNext Poly(A) mRNA Magnetic Isolation Module (New England Biolabs, Ipswich, Massachusetts) followed by library preparation using the NEBNext Ultra II Directional RNA Library Prep Kit for Illumina (New England Biolabs). Sequencing was performed using a NextSeq 500 (Illumina, San Diego, California) at a target depth of 25 million 2 × 75 bp paired end reads per sample. Reads were quasi-mapped to the mouse genome (mm10 assembly) and quantified at the transcript level using Salmon v0.11.2[86] followed by summary to the gene level using tximport v1.10.1[87]. Differential gene expression was performed with DESeq2 v1.22.2 [88]. Gene set enrichment analysis was performed with the GO Biological Processes[89,90] gene sets fromMSigDB[32] using FGSEA v1.8.0[91] run with 100,000 permutations. 4-group intersections were visualized with UpSetR plots [92]. All other plots were generated in figures 2 – 5 were generated using ggplot2 unless otherwise specified [93].

### Stereotactic FUS mediated BBBD

Sonications using the stereotactic frame were performed using a 1-MHz spherical-face single-element FUS transducer with a diameter of 4.5 cm (Olympus). FUS (0.4 MPa or 0.8 MPa; 120 s, 10-ms bursts, 0.5-Hz burst rate) was targeted to the right striatum. The 6-dB acoustic beamwidths along the axial and transverse directions are 15 mm and 4 mm, respectively. The waveform pulsing was driven by a waveform generator (AFG310; Tektronix) and amplified using a 55-dB RF power amplifier (ENI 3100LA; Electronic Navigation Industries).

Once anesthetized, a tail-vein catheter was inserted to permit i.v. injections of MBs and Evans Blue. The heads of the mice were shaved and depilated, and the animals were then positioned prone in a stereotactic frame (Stoelting). The mouse heads were ultrasonically coupled to the FUS transducer with ultrasound gel and degassed water and positioned such that the ultrasound focus was localized to the right striatum. Mice received an i.v. injection of the MBs (1 × 10^5^ MBs/g b.w.) and Evans Blue, followed by 0.1 mL of 2% heparinized saline to clear the catheter. Sonication began immediately after clearance of the catheter. In contrast to the MR-guided experiments, which targeted four spots, only one location was targeted in these studies due to the increased focal region of the transducer (4 mm in the transverse direction, relative to 1 mm for the transducer in the MR-compatible system).

### Histological Processing and Analysis

60 minutes after Evans Blue injection, mice were euthanized via an overdose of pentobarbital sodium and phenytoin sodium. A macroscopic image was taken immediately after whole brain harvest. Brains were then placed in 10% NBF, embedded in paraffin, and sectioned 400 μm apart. H&E stained sections were imaged with 4x and 20x objectives on an Axioskop light microscope (Zeiss, Germany) equipped with a PROGRES GRYPAX microscope camera (Jenoptik, Germany). 10 20x images from the region of the right striatum with maximal Evans Blue extravasation were taken per section and 2 – 6 sections were imaged per brain. A researcher blinded to treatment condition assigned a score of 0 (none), 1 (mild), 2 (moderate), or 3 (severe) to each 20x image for RBC extravasation and vacuolation using a custom MATLAB (MathWorks) script.

## Acknowledgements

Supported by National Institutes of Health Grants R01EB020147, R01CA197111, R01NS111102, and R21EB024323 to R.J.P. and R01EB023055 to A.L.K. A.S.M. was supported by National Institutes of Health Training Grant T32LM012416. C.M.G. was supported by American Heart Association Fellowship 18PRE34030022 and a UVA School of Medicine Wagner Fellowship. N.D.S. was supported by a National Cancer Institute F99/K00 Predoctoral to Postdoctoral Fellow Transition Award (F99CA234954), NSF Graduate Research Fellowship, and a UVA School of Medicine Wagner Fellowship. We thank the UVa Genome and Technology Core for their assistance with sample processing.

## Conflict of Interest

The authors declare no conflict of interest.

## Data Availability

Bulk RNA sequencing data have been deposited in the Gene Expression Omnibus database (https://www.ncbi.nlm.nih.gov/geo/query/acc.cgi?acc=GSE152171). All remaining data generated or analyzed during this study are included in this article.

## Author Contributions

Conceptualization - A.S.M., C.M.G., N.D.S., E.A.T., and R.J.P.; Methodology - A.S.M., C.M.G., N.D.S., E.A.T., W.J.G., A.L.K., G.W.M., and R.J.P.; Investigation - A.S.M., C.M.G., N.D.S., E.A.T., W.J.G., A.L.K., G.W.M., and R.J.P.; Formal Analysis - A.S.M., C.M.G., N.D.S., E.A.T., and R.J.P.; Writing – Original Draft Preparation, A.S.M. and R.J.P.; Writing – Review & Editing, A.S.M., C.M.G., N.D.S., E.A.T., W.J.G., A.L.K., G.W.M., and R.J.P.; Supervision, G.W.M. and R.J.P.; Funding Acquisition - A.S.M. and R.J.P.

## Notes

### Competing Interest Statement

The authors have declared no competing interest.

