## Supplemental Table for "Transcriptomic Response of Brain Tissue to Focused Ultrasound-Mediated Blood-Brain Barrier Disruption Depends Strongly on Anesthesia"

**This PDF file includes:**

Table S1
SI References

**Table S1**

| | Reference | Disease model | Therapeutic | Anesthesia details |
|---|---|---|---|---|
| ISOFLURANE | 1 | Glioma | Polymeric nanoparticles | Isoflurane |
| | 2 | Naive | N/A | 2-3% Isoflurane |
| | 3 | Naive | Dextrans | 1.5% Isoflurane |
| | 4 | Naive | Dextrans | 2-3% Isoflurane |
| | 5 | Alzheimer's disease | N/A | 1-2% Isoflurane |
| | 6 | Naive | Nanoparticles | 2-2.5% Isoflurane |
| | 7 | Huntington's disease | GDNF plasmid | 2% Isoflurane |
| | 8 | Naive | N/A | 2% Isoflurane (vol/vol) in MA |
| | 9 | Glioma | Carboplatin | 2-3% Isoflurane in air |
| | 10 | Naive | Polymeric nanoparticles | Induction: 2% Isoflurane in 20% O2/78% MA, Maintenance: 2% Isoflurane in MA |
| | 11 | Naive | Gad-based nanoparticles | 1.5-2% Isoflurane "in a mixture of air and oxygen" |
| | 12 | Naive | Alpha-synuclein gene | Induction: 3% Isoflurane in O2, Maintenance: 2% Isoflurane in MA |
| | 13 | Naive | NA | Induction: 5% Isoflurane (1 L/min) in O2, Maintenance: 1-2% Isoflurane (1 L/min) in MA |
| | 14 | Naive | rAAV | 2% Isoflurane (vol/vol) in 50% O2/50% MA |
| | 15 | Naive | NA | 1–3.5% Isoflurane (vol/vol) in O2 |
| | 16 | GBM | Doxirubicin | Isoflurane in O2 |
| | 17 | Naive | IP BRDU after FUS, nothing coinjected | 1-3.5% Isoflurane (vol/vol) in O2 |
| | 18 | Naive | Gold nanoclusters | 1-2% Isoflurane (vol/vol) in O2 |
| | 19 | Naive | N/A | 2% Isoflurane (vol/vol) in O2 |
| | 20 | Naive | N/A | Induction: 3% Isoflurane in O2, Maintenance: 2% Isoflurane in O2 |
| | 21 | Naive | Polymeric nanoparticles | 1.5-2% Isoflurane in O2 |
| | 22 | Naive | N/A | 2-3% Isoflurane in O2 |
| | 23 | Naive | N/A | 1.25% Isoflurane + 0.8 ppm O2 |
| | 24 | GBM | IL-12 | 2% Isoflurane (0.8 L/min) in O2 |
| | 25 | GBM | Gene and folate conjugated MB | 1% Isoflurane (1 L/min) in O2 |
| | 26 | Parkinson's disease | Intranasal BDNF | 1-2% Isoflurane in O2 |
| | 27 | Naive | N/A | Induction: 2% Isoflurane in O2, Maintenance: 0.5% Isoflurane in O2 |
| | 28 | Naive | N/A | 1-3.5% Isoflurane in O2 |
| KETAMINE + $\alpha_2$ AGONIST | 29 | Parkinson's disease | GDNF transgene | Ketamine (40 mg/kg) + Dexmedetomidine (0.2 mg/kg) |
| | 30 | Naive | Polymeric nanoparticles | Ketamine (40 mg/kg) + Dexmedetomidine (0.2 mg/kg) |
| | 31 | Glioma | Polymeric nanoparticles | Ketamine (40 mg/kg) + Dexmedetomidine (0.2 mg/kg) |
| | 32 | Naive | Polymeric nanoparticles | Ketamine (40 mg/kg) + Dexmedetomidine (0.2 mg/kg) |
| | 33 | Naive | Reporter gene nanoparticles | Ketamine (40 mg/kg) and Dexmedetomidine (0.2 mg/kg) |
| | 34 | Naive | Herceptin (trastuzumab) | Ketamine (70 mg/kg) + Xylazine(10 mg/kg) |
| | 35 | Alzheimer's disease | Endogenous IgG | Ketamine (150 mg/kg) + Xylazine (10 mg/kg) |
| | 36 | Naive | AAV for GFP | Ketamine (90 mg/kg) + Xylazine (4 mg/kg) |
| | 37 | Naive | Evans Blue | Ketamine (75 mg/kg) + Xylazine (4 mg/kg) |
| | 38 | Naive | N/A | Claims Ketamine (90 mg/kg) and Xylazine (10 mg/kg) |
| | 39 | Metastatic BrCa | Therapeutic antibodies | Ketamine (80 mg/kg) + Xylazine (10 mg/kg) |
| | 40 | Glioma | Liposomal Doxorubicin | Ketamine (80 ml/kg/h) + Xylazine (10 ml/kg/h) |
| | 41 | Metastatic BrCa | Trastuzumab | Ketamine (90 mg/kg) + Xylazine (10 mg/kg) |
| | 42 | Naive | N/A | Ketamine (40–50 mg/kg) + Xylazine (10 mg/kg) |
| | 43 | Alzheimer's disease | N/A | Ketamine (75 mg/kg) + Xylazine (4 mg/kg) |
| | 44 | Naive | Dually labeled liposomes | Ketamine (80 mg/kg/hr) + Xylazine (10 mg/kg/hr) |
| | 45 | Naive | N/A | Ketamine (80 mg/kg) + Xylazine (10 mg/kg) |
| | 46 | Alzeheimer's disease | Anti-tau antibody | Ketamine (100 mg/kg) + Xylazine (10 mg/kg) |
| MISCELLANEOUS | 47 | Naive | N/A | Induction: 3% Isoflurane (vol/vol) in MA, Maintenance: Dexmedetomidine (0.1 mg/kg) |
| | 48 | Naive | N/A | For sedate animals: Ketamine (10 mg/kg) + Atropine (0.02-0.04 mg/kg) to induce, 1-2% Isoflurane/O2 to maintain. For awake animals: Ketamine (5 mg/kg) |
| | 49 | Naive | N/A | Induction: Ketamine (50 mg/kg/h) + Xylazine (5 mg/kg/h), Maintenance: 1-3% Isoflurane (2 L/min) in MA |
| | 50 | Naive | N/A | Zoletil (25 mg/kg) and Rompun (4.6 mg/kg) |
| | 51 | Naive | N/A | Induction: Tiletamine (6 mg/kg) + Xylazine (2.2 mg/kg), Maintenance: Propofol (10 mg/kg/hr) |
| | 52 | Naive | GABA | Induction: Ketamine (3 mg/kg) + Dexmedetomidine (0.015 mg/kg), Maintenance: 1% Isoflurane in O2 |
| | 53 | Naive | Doxorubicin | Zoletil (25 mg/kg) and Rompun (4.6 mg/kg) in saline |
| | 54 | Naive | N/A | Telazol (tiletamine and zolazepam): 2-4 mg/kg |
| | 55 | Metastatic melanoma | Polymeric nanoparticles | 2:1:2:5 mixture of Fentanyl, Medetomidine, Midazolam, and water, (10 ul/g), subcutaneous injection |
| | 56 | GBM | Doxorubicin | gas anesthesia |
| | 57 | GBM | BCNU | Chlorohydrate (30 mg/kg) |
| | 58 | Naive | N/A | Chlorohydrate (30 mg/kg) |
| | 59 | GBM | TMZ | Chlorohydrate (30 mg/kg) |
